# MIDFA: Scalable Bayesian Factor Analysis for Mixed and Incomplete Data

**DOI:** 10.64898/2026.06.10.731315

**Authors:** George Hutchings, Pantelis Samartsidis, Corinne Donnay, Laura Gaetano, Elizabeth Fisher, Thomas E. Nichols, Chris Holmes, Dieter A. Häring, Habib Ganjgahi

## Abstract

Probabilistic latent variable models are a powerful tool for uncovering structure in high-dimensional datasets, particularly in biomedical applications. The increasing availability of large-scale epidemiological studies, such as the UK Biobank, poses important modelling challenges, including mixed data types, high dimensionality, and structured missingness. Existing approaches address some of these issues, but few provide a unified and scalable framework for handling them simultaneously. Here, we propose a scalable Bayesian factor analysis framework designed to address these challenges. Our method combines a semi-parametric Gaussian copula model with a continuous spike-and-slab prior to induce sparse and interpretable factor loadings. The number of latent dimensions is learned nonparametrically from the data using an Indian buffet process prior. For model fitting, we develop an expectation–maximisation algorithm that naturally accommodates missing data. We validate the proposed method through comprehensive simulation studies. In addition, we showcase the proposed model using the Novartis–Oxford Multiple Sclerosis dataset in two ways. First, we identify latent dimensions shared across MS clinical and neuroimaging variables, characterising disease structure while demonstrating the model’s ability to handle multiple data types and structured missingness. Second, we use the model for dimensionality reduction of structural MRI data, extracting features for downstream analysis that go beyond traditional whole-brain summary statistics. In those applications, our method identifies sparse latent structures and provides insights beyond those obtained from traditional approaches.

## 1 Introduction

The rise of large scale epidemiological studies, such as the UK Biobank (Sudlow et al., 2015), the Novartis-Oxford Multiple Sclerosis (NO.MS) dataset (Dahlke et al., 2021), the Adolescent Brain Cognitive Development (ABCD) study (Casey et al., 2018), and the Human Connectome Project (HCP) (Van Essen et al., 2012), is creating a new paradigm for biomedical research. Dimensionality reduction is an important technique for the analysis of such datasets. By providing a more interpretable, low dimensional representation of the data’s structure, this compression enables a wide range of benefits, including the visualisation of high dimensional data, the discovery of latent structure to uncover biological patterns, outlier removal for data quality control, data fusion to integrate information from mixed data types, and feature extraction to enable downstream modelling, a task that is often computationally infeasible with the full dataset.

Despite their potential, these datasets possess several intrinsic characteristics that make dimensionality reduction challenging. First, they involve mixed data types, containing a combination of continuous, binary, and ordinal variables. Second, they are often complicated by structured missingness, a non random pattern of missing data inherent to how the studies were conducted. Third, the true number of latent dimensions is not known in advance. Fourth, the scale of the data is large, both in its dimensionality, particularly with the inclusion of imaging data, and its number of individuals. Finally, although any compressed data can be valuable for downstream prediction, gaining direct scientific insight requires that the latent dimensions themselves are interpretable.

Many methods for dimensionality reduction exist, but they are often poorly equipped to handle these challenges concurrently. Classical techniques like principal component analysis (PCA) (Pearson, 1901), Factor Analysis (FA) (Spearman, 1961), Independent Component Analysis (ICA) (Comon, 1994), and Nonnegative Matrix Factorisation (NMF) (Kim & Park, 2008) are effective for single type, fully observed data but do not handle mixed data type variables, accommodate missing values, or infer the number of latent dimensions. More advanced methods have been developed to address some subsets of these limitations, for example, by providing frameworks for handling mixed data (Murray et al., 2013) or inferring sparse, interpretable structure (Li et al., 2023; Ročková & George, 2016), but they fall short of providing a complete solution.

A significant challenge is the integration of mixed data types. A common strategy, as used by Feldman and Kowal (2022), Murray et al. (2013), and Quinn (2004), is to combine a factor analysis model with a Gaussian copula (Sklar, 1959). The factor model provides the mechanism for dimensionality reduction, decomposing the observed data into factor scores, which summarise each individual on a smaller set of latent variables, and factor loadings, which describe how each observed variable relates to these latent dimensions. The Gaussian copula, in turn, provides the link for handling mixed data types, assuming that all variables, regardless of their observed type, arise from these underlying continuous latent variables, which have a joint Gaussian distribution. For ordinal data, standard approaches estimate category cutpoints as parameters, typically via Markov Chain Monte Carlo (MCMC), which can harm scalability (Quinn, 2004). To overcome the difficulties of estimating these cutpoints, Feldman and Kowal (2022) and Murray et al. (2013) use semiparametric Gaussian copula models. This approach leverages the extended rank likelihood (ERL) proposed by D. Hoff (2007), which considers only the ranks of the observed data, thereby avoiding the need to explicitly estimate the cutpoints. Alternative approaches, such as Multiple Correspondence Analysis (MCA) (Le Roux & Rouanet, 2010), extend principal component analysis to categorical data but do not have the capability to handle missing data, infer the number of latent dimensions, or provide interpretable solutions.

The presence of missing values is another major hurdle. Common ad hoc strategies like complete case analysis, where only fully observed individuals are analysed, are infeasible, as the large volume of missing data would lead to a loss of power and bias (Sterne et al., 2009). An alternative is imputation. However, single imputation methods are statistically invalid as they fail to account for imputation uncertainty and may also lead to bias (Sterne et al., 2009). While more principled methods like Multiple Imputation (MI) (Rubin, 1987) correctly account for this uncertainty, they face two major obstacles in this context. First, the sheer volume and structured nature of the missingness (e.g. entire blocks of variables unobserved for certain subgroups) make specifying a reliable imputation model challenging and can thus violate the assumptions of standard MI algorithms. Second, even if a suitable model could be specified, the requirement to fit the analysis model multiple times can be computationally prohibitive for the large scale datasets.

Determining the true number of latent dimensions, *K*, is a critical model selection problem. Most standard methods require *K* to be fixed in advance. A common strategy is therefore to fit the model repeatedly over a range of candidate values for the number of latent dimensions, *K*. The optimal value is then selected using an appropriate model selection criterion, such as the Akaike Information Criterion (AIC), the Bayesian Information Criterion (BIC), or an evaluation of the proportion of variance explained (Hastie et al., 2009). However, this selection procedure fails to propagate uncertainty about *K*: once a value is chosen, it is treated as fixed and known, yielding over-optimistic uncertainty quantification. This process is computationally expensive for large datasets and can be subjective, yet remains a necessary step for many frameworks.

Another challenge is non-identifiability, classical latent variable models are identifiable only up to orthogonal transformations of the latent variables and loading matrix, so many observationally equivalent solutions exist. Principal component analysis enforces orthogonality of the components, yielding a unique solution, up to sign and ordering when eigenvalues are distinct, whereas exploratory factor analysis typically applies a post hoc rotation towards a sparse solution. A sparse loading matrix can resolve the rotational ambiguity and aids interpretation, as each latent dimension is then defined by a small, distinct subset of features. However, standard post hoc approaches such as Varimax rotation (Kaiser, 1958) involve an arbitrary choice of rotational criterion and still produce dense loadings, requiring a further thresholding step to achieve exact zeros. Another approach to enforce a unique solution is to impose constraints, such as forcing the loading matrix to be upper triangular (Murray et al., 2013). While this guarantees identifiability, the resulting structure is an artefact of the constraint itself and is not necessarily scientifically meaningful.

A more principled, model based approach is to induce sparsity directly through the prior distribution of the factor loadings, which simultaneously addresses identifiability and enhances interpretability. While common regularisation priors (Hastie et al., 2009) based on *L*_1_ (LASSO) or *L*_2_ (Ridge) norms are computationally efficient, they do not produce exact zeros and introduce shrinkage bias by shrinking large, non zero coefficients towards zero.

An alternative is the spike and slab prior (Mitchell & Beauchamp, 1988), which models each loading as a draw from a two component mixture. A high concentration spike distribution (centred at zero) captures negligible feature contributions, while a diffuse slab distribution accommodates significant, non zero loadings. This approach can distinguish between important and unimportant features, shrinking only the latter. However, the choice of spike and slab formulation carries computational trade offs. Discrete versions that use a point mass at zero as the spike offer the clearest formulation but are often computationally intractable in high dimensions. Continuous relaxations have been proposed (George & McCulloch, 1993) as scalable alternatives, with similar approaches seen in the dimensionality reduction methods of Li et al. (2023) and Ročková and George (2016), which have not been extended to mixed-type data.

Finally, the scale of modern biomedical datasets, composed of data from thousands of individuals with tens of thousands of features, poses a significant computational barrier. While classical methods like PCA are usually computationally tractable due to their simplicity, this tractability comes at the cost of the flexibility needed to handle the complex data characteristics discussed previously. In contrast, many Bayesian models are limited in their practical application. They either rely on full MCMC sampling, which is computationally infeasible for datasets of this size (Murray et al., 2013; Quinn, 2004), or, lack closed form analytical solutions, and so must resort to computationally expensive optimisation routines (Li et al., 2023; Ročková & George, 2016). This creates a critical need for methods that can bridge this gap, offering both the flexibility to model complex data and the computational efficiency to operate at scale. Consequently, a unified framework that can concurrently handle all these characteristics at scale remains a significant gap in the field.

Our primary contribution is a unified and scalable Bayesian factor analysis framework, which we term MIDFA (Mixed Incomplete Data Factor Analysis), designed to address several concurrent challenges presented by modern biomedical data. Specifically, we introduce: (1) a single, coherent model, detailed in Section 2, that integrates a semiparametric Gaussian copula for mixed data, a nonparametric Indian Buffet Process prior to infer the latent dimensionality, and a continuous spike and slab prior to learn a sparse factor structure; (2) a scalable inference scheme using an Expectation Maximisation algorithm, presented in Section 2.1, where the choice of priors allows for efficient, closed form updates that avoid the computational burden of full MCMC; and provide (3) a comprehensive validation of our method, with results presented in Section 3, including application to data with a high degree of structured missingness and high dimensional binary-valued neuroimaging data.

## 2 Method

A key challenge in modelling real world datasets is to capture the dependence between variables without imposing restrictive assumptions on their marginal distributions. The Gaussian copula Factor Model (GCFM; Murray et al. (2013)) achieves this by decoupling the marginal distributions from the dependence structure through the use of a copula representation.

Let *Y*_*n*_ = (*Y*_*n*1_, …, *Y*_*nD*_)^⊤^ denote the *D* observed variables for individual *n* (*n* = 1, …, *N*) and *N* is the number of individuals, where each *Y*_*nd*_, *d* = 1, …, *D*, may be continuous, ordinal, or binary. According to Sklar’s theorem (Sklar, 1959), any joint cumulative distribution function (CDF) *F*(*Y*_*n*_) can be expressed in terms of its univariate marginal CDFs *F*_*d*_ and a copula function *C* that captures their dependence. A *D*-dimensional copula *C* : [0, 1]^*D*^ → [0, 1] is a multivariate distribution function with uniform [0, 1] marginals that encodes the dependence structure among the variables, irrespective of their marginal distributions.

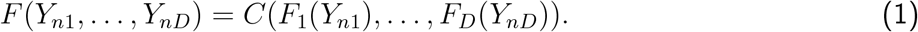

The Gaussian copula specifies this dependence through a correlation matrix *R*, such that

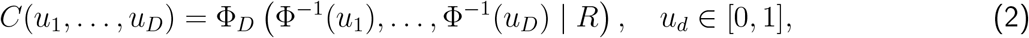

where Φ_*D*_(· | *R*) denotes the *D*-dimensional multivariate normal distribution with zero mean and correlation matrix *R*, and Φ^−1^ is the univariate standard normal quantile function.

Hence, the joint distribution can be written as:

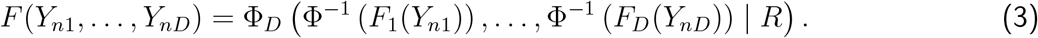

By Sklar’s theorem (Sklar, 1959), if all *F*_*d*_ are continuous then the copula is unique; if any margins are discrete it is uniquely determined on Ran(*F*_1_) × · · · × Ran(*F*_*D*_), where Ran denotes the range of *F*_*d*_.

In line with Murray et al. (2013), a copula factor model can be described as follows. Let Σ be a covariance matrix with correlation matrix *R* and draw *Z*_*n*_ ~ 𝒩 (0, Σ); with 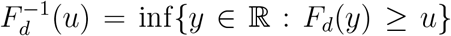 define 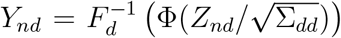 for *d* = 1, …, *D*. Then *Y*_*n*_ has a Gaussian copula with correlation matrix *R* and univariate marginals *F*_*d*_. This representation can be used to generalise the Gaussian factor model to the Gaussian copula factor model.

The GCFM introduces a latent Gaussian representation to impose a factor structure on Σ. Specifically, each observation *Y*_*n*_ is associated with a latent Gaussian vector *Z*_*n*_ = (*Z*_*n*1_, …, *Z*_*nD*_)^⊤^ satisfying

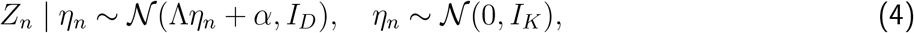

where Λ ∈ ℝ^*D*×*K*^ is the factor loading matrix, with Λ_*d*_ denoting its *d*th row and Λ_*dk*_ its (*d, k*) element, *η*_*n*_ ∈ ℝ^*K*^ are the *K*-dimensional factor scores (treated as column vectors), *α* ∈ ℝ^*D*^ are intercepts controlling feature-specific means, and *I*_*D*_ is the identity matrix of size *D* × *D*. The correlation matrix *R* of the latent Gaussian variables is therefore constrained to follow a low rank structure.

The observed variables *Y*_*n*_ are linked to the latent Gaussian variables *Z*_*n*_ through their feature specific marginal CDFs. Because each latent dimension *Z*_*nd*_ has marginal variance 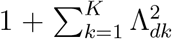, it must be standardised to unit variance before applying the Gaussian copula transformation. The mapping to the observed scale is therefore

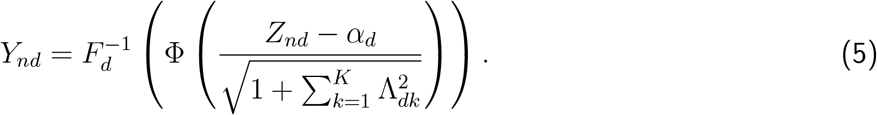

This scaling ensures that each transformed latent variable has a standard normal marginal, maintaining a valid copula representation.

This separation between the dependence structure (governed by the copula through *Z*_*n*_) and the marginals (through *F*_*d*_) allows the model to flexibly handle mixed data types.

In practice, the marginal distributions *F*_*d*_ are typically unknown and treated as nuisance parameters. To avoid parametric assumptions and improve robustness, we adopt a semiparametric approach using the ERL. This likelihood exploits the fact that the transformation of eq. (5) is monotonic, meaning that the observed data impose an ordering constraint on the latent variables. Denote by 𝒟 (*Y*) the set of all latent matrices *Z* = [*Z*_1_, …, *Z*_*N*_]^⊤^ ∈ ℝ^*N*×*D*^ consistent with the observed ranks of *Y* = [*Y*_1_, …, *Y*_*N*_]^⊤^ ∈ ℝ^*N*×*D*^:

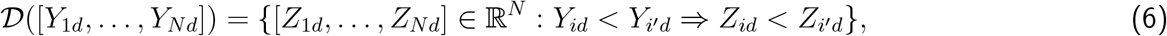

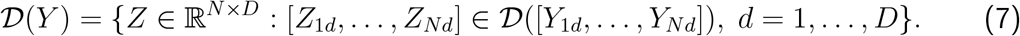

The semiparametric likelihood is then defined as

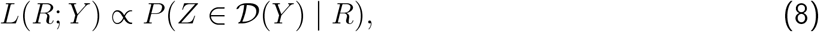

which depends only on the copula correlation matrix *R*. This approach preserves most of the information relevant to dependence, while avoiding explicit modelling of the marginal distributions, and is used in place of the full likelihood throughout inference.

The Gaussian copula Factor Model provides a flexible framework for capturing complex, non Gaussian dependence structures through a latent Gaussian representation, while remaining robust to the form of the marginal distributions. Murray et al. (2013) additionally prove strong posterior consistency for the correlation matrix under the ERL, offering theoretical assurance for its application to mixed data. Computationally, this formulation is also advantageous, as the constraints defined by 𝒟(*Y*) lead to a straightforward Gibbs sampling step where each latent variable *Z*_*nd*_ is drawn from a truncated normal distribution.

When all observed variables are continuous, a two step alternative may be used (Murray et al., 2013). Each observation is transformed via 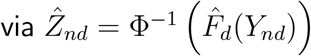 using the empirical CDF 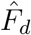, and the ‘pseudo data’, 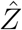, is then treated as fixed when fitting the factor model on the latent Gaussian scale. This approach avoids the rank likelihood approximation and can reduce computation but requires fully continuous data.

While binary data can be treated as a special case of ordinal data within the ERL framework, this imposes a potentially meaningless ordering on the outcomes (e.g. that 1 > 0 in binary). A more direct and conceptually natural approach is a probit formulation (Albert & Chib, 1993), which avoids this issue. As argued by Feldman and Kowal (2022), this is more appropriate for binary and unordered categorical data. Under this construction, an observed binary outcome *Y*_*nd*_ ∈ {0, 1} is assumed to arise from its corresponding latent variable *Z*_*nd*_ crossing a zero threshold:

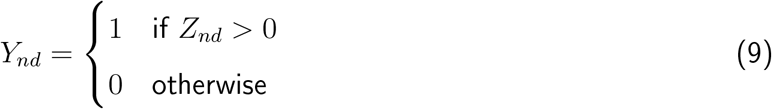

This imposes fixed, known truncation bounds on *Z*_*nd*_ ((−∞, 0] for an observation of 0 and (0, ∞) for an observation of 1), which simplifies the Gibbs sampling step compared to the data dependent bounds of the ERL, making it computationally advantageous. This probit model is a special case of the more general extended rank probit likelihood proposed by Feldman and Kowal (2022) for handling unordered categorical data. This formulation requires a feature specific intercept, *α*_*d*_, to allow the model to capture the baseline marginal probability of the outcome. For unordered categorical features with *K*_*d*_ levels, replace *Y*_*nd*_ by a one-hot indicator block 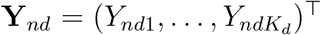 with *Y*_*ndk*_ = 𝕀{*Y*_*nd*_ = *k*}. Augment *Y*_*n*_ by substituting **Y**_*nd*_ for *Y*_*nd*_ and fit the same probit factor model to the enlarged vector (Feldman & Kowal, 2022).

The factor scores and loading matrix in classical factor analysis models are only determined up to an orthogonal transformation, which results in the existence of infinitely many equivalent solutions. Among many solutions to achieve identifiability, we use a sparsity inducing prior on the loading matrix, following the approach of Ročková and George (2016). Specifically, we use a continuous spike and slab prior (George & McCulloch, 1993). This prior models each loading element, Λ_*dk*_, as being drawn from a two component Gaussian mixture. A spike distribution, 𝒩(0, *v*_0_), with a very small variance, shrinks the negligible loadings towards zero. Conversely, a diffuse slab distribution, 𝒩(0, *v*_1_), with a large variance, accommodates substantial non zero loadings. The allocation to either component is governed by a binary indicator variable, *w*_*dk*_ ∈ {0, 1}, such that:

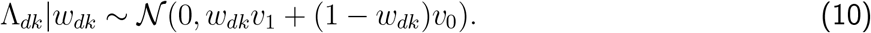

for *d* = 1, …, *D* and *k* = 1, …, *K*, where 0 *< v*_0_ ≪ *v*_1_. In practice (Ročková & George, 2016), this selection is performed by examining the Posterior Inclusion Probability (PIP) for each loading, calculated as the posterior probability *P*(*w*_*dk*_ = 1|data). A loading is included in the model if its PIP exceeds a predefined threshold (e.g. 0.5), signifying it is more likely to belong to the slab.

To determine the latent dimensionality, we adopt a nonparametric Bayesian approach that allows the model complexity to be learned directly from the data. We place an Indian Buffet Process (IBP) prior on the binary indicator matrix *W* ∈ {0, 1}^*D*×*K*^, with elements *w*_*dk*_, that governs the spike and slab allocations (Ročková & George, 2016). The IBP defines a prior over binary matrices with a potentially infinite number of columns, allowing the number of active latent dimensions to be inferred directly from the data (Ghahramani & Griffiths, 2005). For computational tractability within our framework, we employ the stick breaking construction of the IBP (Teh et al., 2007). This process generates a sequence of decreasing, dimension specific inclusion probabilities, *θ*_*k*_ for *k* = 1, … *K*:

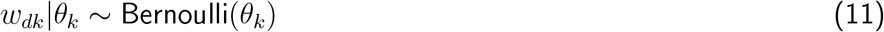

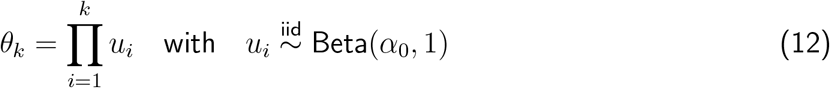

This construction imposes an ordering that makes latent dimensions with higher indices progressively less likely to be active. This mechanism encourages later ones to be inactive, allowing the number of active latent dimensions to be inferred from the data. The hyperparameter *α*_0_ controls the expected number of active latent dimensions a priori. This ordering also serves as a soft identifiability constraint against the permutational invariance of the factor model. For practical computation, the number of latent dimensions is truncated at a large value *K*, which is chosen to be sufficiently high so as not to constrain the posterior distribution.

### 2.1 Model Fitting

We estimate the model parameters Θ = {Λ, *α, θ*} and latent variables *Ƶ* = {*η, Z, w*} by maximising the joint posterior *p*(Θ, *Ƶ* | *Y*), where for notational convenience we collect the latent factor scores into the matrix *η* = (*η*_1_, …, *η*_*N*_)^⊤^ ∈ ℝ^*N*×*K*^. Direct optimisation is intractable. We therefore employ an Expectation Maximisation (EM) algorithm (Dempster et al., 1977), which iteratively increases a lower bound on the marginal log posterior. Let Θ^(*t*)^ denote the parameter estimates at iteration *t*. The EM algorithm iteratively maximises the expected log augmented posterior, the *Q*-function,

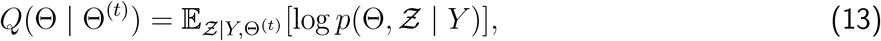

where the expectation is taken over the conditional distribution of the latent variables given the observed data and the current parameter estimates. The algorithm alternates between computing the expectation in eq. (13), the E-step, and maximising *Q*(Θ | Θ^(*t*)^) with respect to Θ to obtain Θ^(*t*+1)^, the M-step. This guarantees a non decreasing sequence of posterior values until convergence to a local mode.

For our hierarchical Gaussian copula model, the augmented posterior satisfies

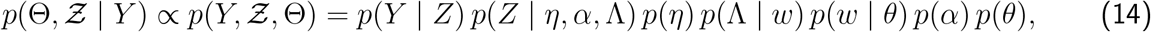

and hence the *Q*-function separates additively into *Q*(Θ | Θ^(*t*)^) = *Q*_Λ,*α*_(Λ, *α* | Θ^(*t*)^) +*Q*_*θ*_(*θ* | Θ^(*t*)^) where,

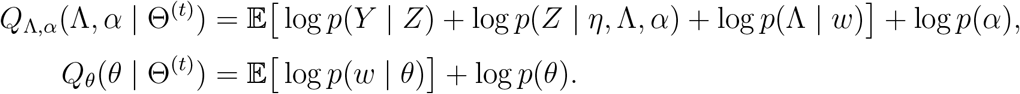

Therefore, the updates naturally decouple into two smaller optimisation problems: updating *θ* requires maximising *Q*_*θ*_(*θ* | Θ^(*t*)^), while updating Λ and *α* requires maximising *Q*_Λ,*α*_(Λ, *α* | Θ^(*t*)^).

#### E-Step

In the E-step we require the conditional expectations of the latent variables *η, Z* and the inclusion indicators *w* given the observed data and the current parameter estimates Θ^(*t*)^. The hierarchical structure of the model allows for a useful separation in the conditional distribution: the latent variables (*η, Z*) and the inclusion indicators *w* become conditionally independent once we condition on *Y* and Θ^(*t*)^. This enables us to compute their expectations efficiently in two parts. First, we evaluate the expectations of (*η, Z*), and second, we evaluate the expectations of *w*, with both sets of expectations taken conditional on the data and Θ^(*t*)^. This decomposition simplifies the E step, and the resulting expectations feed directly into the M step.

The expectations in *Q* that involve *η* and *Z* are intractable, so we approximate them using Monte Carlo integration with *M* samples 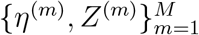 drawn from their full conditional distributions via a Gibbs sampler. Following Ročková and George (2016), we run only a few Gibbs iterations per EM step rather than a full chain. The full conditionals are as follows. For the latent Gaussian variables:

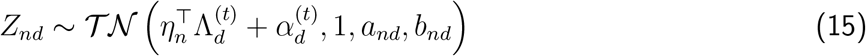

For *n* = 1, … *N, d* = 1, … *D* where *a*_*nd*_, *b*_*nd*_ are the cutpoints induced by the ERL, *a*_*nd*_ = max({*Z*_*i*_*′*_*d*_ : *Y*_*i*_*′*_*d*_ *< Y*_*nd*_}∪{−∞}), *b*_*nd*_ = min({*Z*_*i*_*′*_*d*_ : *Y*_*i*_*′*_*d*_ *> Y*_*nd*_}∪{+∞}). Here 𝒯𝒩 (*µ, σ*^2^, *a, b*) denotes the normal distribution 𝒩 (*µ, σ*^2^) truncated to the interval [*a, b*]. For missing observations 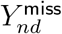, the corresponding latent Gaussian variable 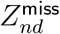 is sampled from its unconditional normal distribution

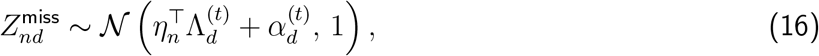

thus providing a natural, fully probabilistic treatment of missing data.

The factor scores are sampled from

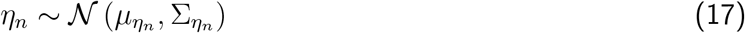

where 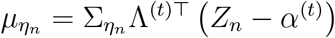 and 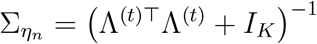.

For the binary inclusion indicators *w*_*dk*_, the expectations are computed in closed form:

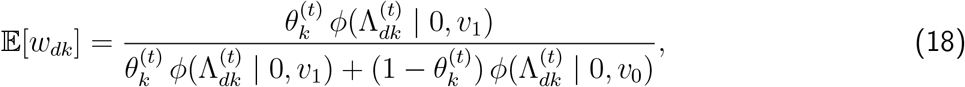

where *ϕ*(· | 0, *v*) denotes the univariate Gaussian density with mean zero and variance *v*.

#### M-step

The maximisation of the approximate *Q*-function with respect to the parameters separates naturally across Λ, *α*, and *θ*. Throughout this section, 𝔼[·] denotes expectation under the E-step posterior *p*(*Ƶ* | *Y*, Θ^(*t*)^). Let *Z*_•*d*_ = (*Z*_1*d*_, …, *Z*_*Nd*_)^⊤^ denote the *d*th column of *Z* and 1_*N*_ the vector of ones of length *N*. We set

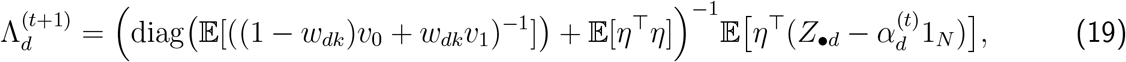

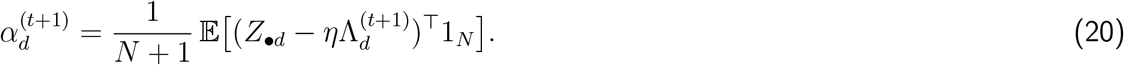

The expectations with respect to (*Z, η*) in eqs. (19) and (20) are intractable, so we approximate them by averaging over the *M* Gibbs samples 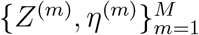 drawn in the E-step. The expectation with respect to *w* in eq. (19) is available in closed form from eq. (18), and the diagonal term simplifies accordingly (see Supplementary Material). The approximate updates are

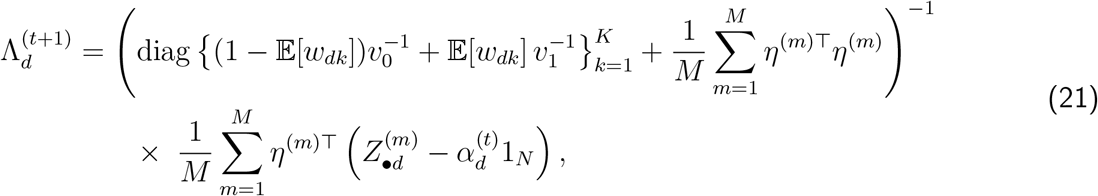

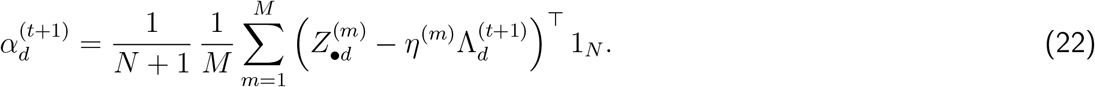

The stick breaking parameters *θ* of the Indian Buffet Process are updated by maximising

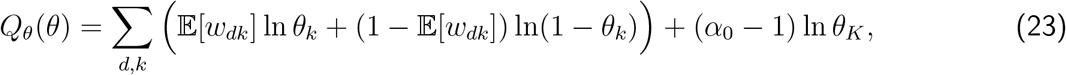

subject to the constraint 1 > *θ*_1_ > *θ*_2_ > · · · > *θ*_*K*_ > 0. This constrained optimisation is performed numerically.

Combining these steps yields the procedure summarised in Algorithm 1, with full derivations given in the Supplementary Material.

##### Algorithm 1

Monte Carlo EM algorithm for hierarchical Gaussian copula factor analysis.

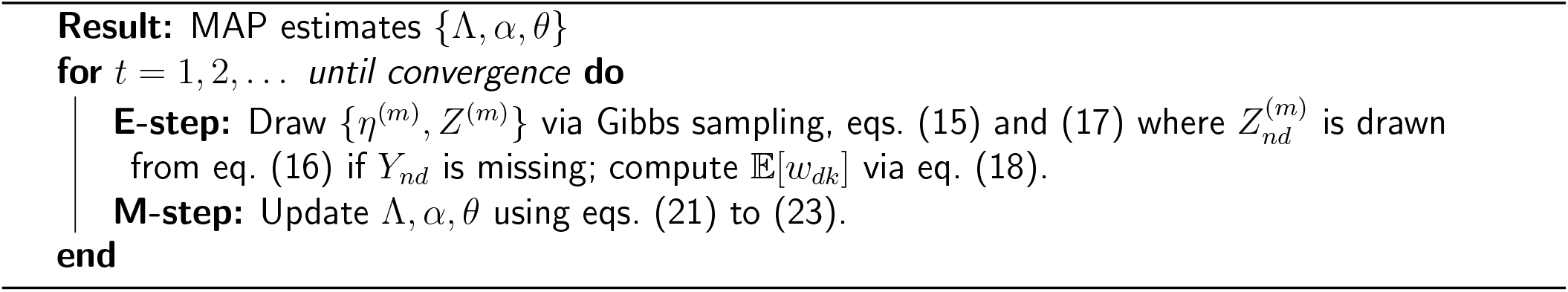

Following convergence, the estimated number of latent dimensions 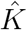 is the number of columns of Λ for which at least one loading has posterior inclusion probability exceeding 0.5; columns with no active loadings are discarded.

#### Dynamic posterior exploration

A challenge in fitting our model is that the EM algorithm is only guaranteed to converge to a local optimum of the posterior distribution, making its performance sensitive to initialisation on the complex and multimodal posterior landscape. To mitigate this issue and enhance the search for the posterior mode of highest density, we use a dynamic posterior exploration strategy (Li et al., 2023; Ročková & George, 2016), a technique adapted from the deterministic annealing literature. Sequentially refitting the model along a user-specified cooling schedule of decreasing spike variance *v*_0_, with each stage warm-started from the previous, creates a solution path that guides the algorithm toward modes of higher posterior density. The procedure is summarised in Algorithm 2. Between successive stages, a varimax rotation (Kaiser, 1958) may optionally be applied to realign the loadings toward a sparser orientation before the next warm-start.

##### Algorithm 2

Dynamic posterior exploration.

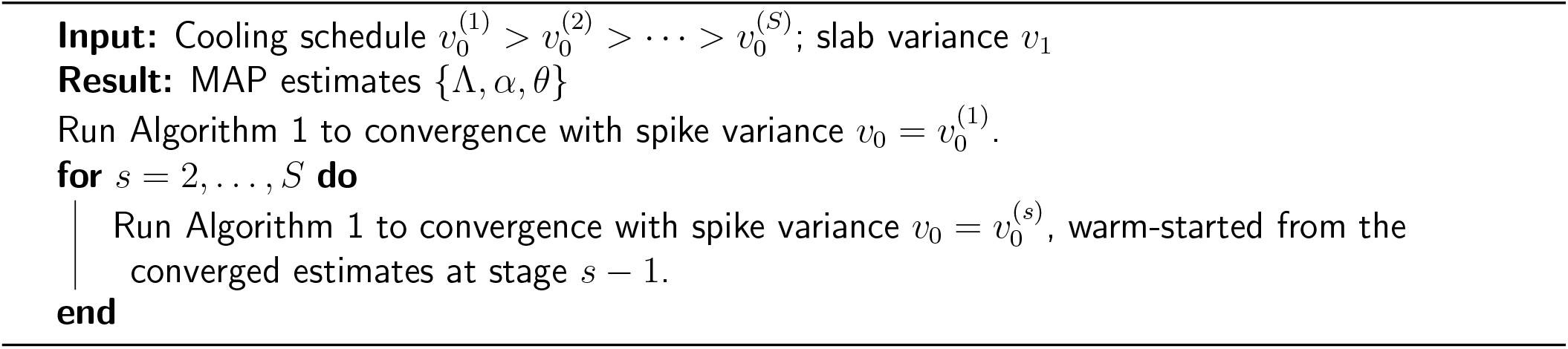

#### Parameter expansion

Although DPE improves the search for good modes, rotational invariance can still slow or misdirect EM, so we next use a parameter expansion that learns a sparse orientation of the loading matrix (Ročková & George, 2016). This expands the parameter space by introducing a positive definite matrix *A* ∈ ℝ^*K*×*K*^. Placing the spike and slab prior on the loadings in this expanded space guides the algorithm toward sparse, well-oriented solutions while leaving the observed data likelihood unchanged.

Let *A*_*L*_ be the Cholesky factor of *A* so that 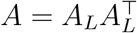. Defining the expanded parameters by 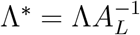 and 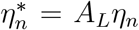, we place the spike and slab prior directly on Λ^∗^. Hence, under the expanded model, 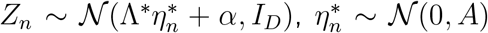 which has the same observed data likelihood and the original model is recovered at *A* = *I*_*K*_. Following Ročková and George (2016), the E-step is unchanged under *A* = *I*_*K*_, while the M-step is augmented to estimate *A*. In particular, we set

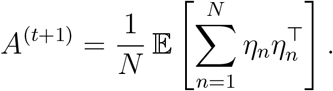

As before, the expectations in this section are approximated by sample averages over the E-step Gibbs samples.

We then apply a rotation using the Cholesky factor 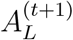:

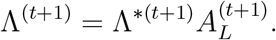

This rotation restores the original parametrisation for the next E-step, so the updated *A* itself is not propagated forward in the E-step. These two modifications implement the automatic rotation to sparsity within EM and improve robustness to random initialisation.

We additionally apply the parameter expansion of Murray et al. (2013) to further aid convergence. We introduce per variable residual variances 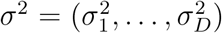, compute the E-step under *σ*^2^ = 1_*D*_, where 1_*D*_ is the vector of ones, length *D*, then set

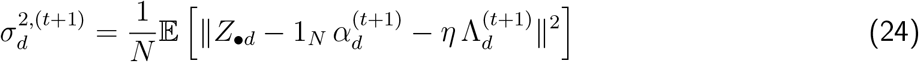

and apply the reduction:

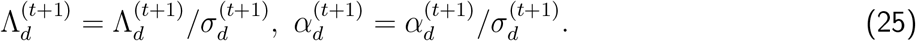

As in Murray et al. (2013) we place independent Gamma priors on the precisions 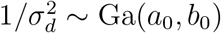 with small *a*_0_, *b*_0_ for a diffuse prior. The updated *σ*^2^ is used only in this reduction and is not carried into the next E-step.

### 2.2 Simulation Studies

To evaluate the performance of our proposed framework, we conduct a series of simulation studies in settings where the ground truth is known. A challenge in such an evaluation is the absence of a single method that concurrently handles mixed data types, accommodates missing values, infers the latent dimensionality, and produces a sparse loading matrix. This precludes a direct comparison against a single, all encompassing benchmark. Therefore, to facilitate fair and rigorous comparisons, we structure our evaluation across three distinct scenarios, each designed to test specific capabilities of our model against relevant, state-of-the-art alternative approaches.

Across all scenarios, data are generated from the underlying factor model using a Gaussian copula to create observations with specified marginal distributions. First, we draw the factor scores *η*_*n*_ ∈ ℝ^*K*^ for each individual *n* = 1, …, *N* from a standard normal distribution, *η*_*n*_ ~ 𝒩 (0, *I*_*K*_). These scores are then projected into the feature space via the loading matrix Λ to produce a latent Gaussian vector, *Z*_*n*_ = Λ*η*_*n*_ + *α* + *ϵ*_*n*_, where *ϵ*_*n*_ ~ 𝒩 (0, *I*_*D*_). The observed data *Y*_*n*_ is then obtained by transforming this latent vector. Each component *Z*_*nd*_ is standardised and mapped to the uniform scale via the standard normal CDF, 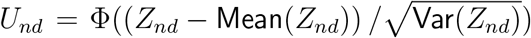. The final observed variable *Y*_*nd*_ is produced by applying the inverse CDF, 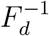, corresponding to the desired data type for that feature. For continuous data, we set the marginal distribution to be Gaussian (*F*_*d*_ = Φ), while for other data types, appropriate marginals are chosen to generate binary or ordinal outcomes. For all simulations, we use a true loading matrix Λ with *D* = 12 features and *K* = 3 latent dimensions, as depicted in Figure 1. This matrix is designed with a sparse structure to provide a test of structural recovery. For all scenarios, the IBP hyperparameter *α*_0_ is set to 2, *N* = 200, and we use *R* = 500 realisations. The IBP stick-breaking representation is truncated at a maximum of *K* = 10 latent dimensions. We do not use parameter expansion or varimax rotation within each DPE stage, though a varimax rotation is applied to the loading matrix between successive stages. The DPE schedule is *v*_0_ = 10^−1^, 10^−2^, 10^−3^, 10^−4^, with *v*_1_ = 10. Within each stage, the E-step is approximated using 100 samples. Convergence is assessed by the maximum absolute change in loading parameters, with a tolerance of 0.016.

**Figure 1:**
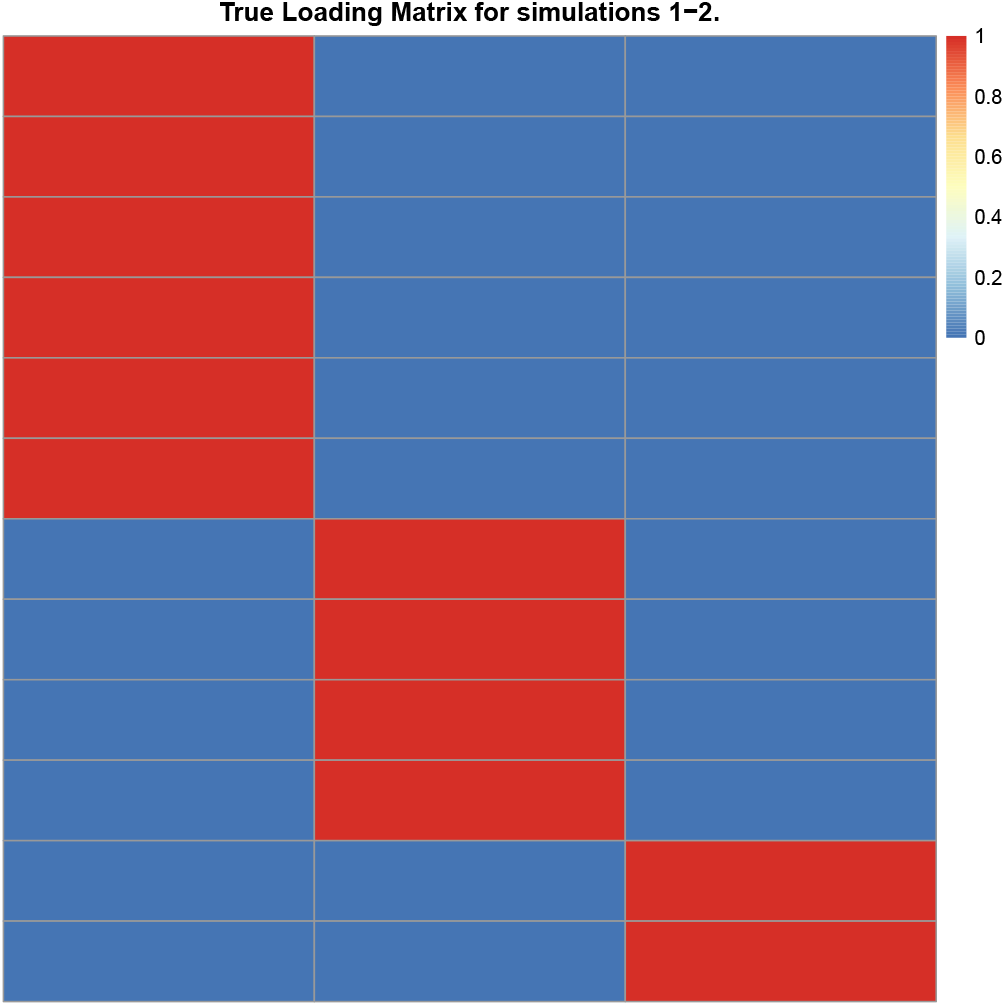
True 12 × 3 loading matrix used in simulation scenarios 1–2.

We assess model performance based on the ability to recover the true loading matrix Λ and its properties. We report three metrics: loading MSE, 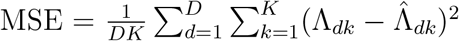; structure recovery, 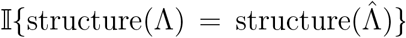, which equals 1 when the sparsity pattern of 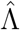 matches that of Λ exactly; and correct 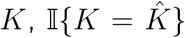, which equals 1 when the estimated number of factors equals the true number.

To address the inherent rotational and sign ambiguity of the loading matrix, we apply a post processing step to all estimated matrices. The true matrix Λ is deliberately constructed such that each column has a unique number of non zero elements. This design allows us to unambiguously permute the columns of any estimated matrix 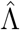 and flip their signs to achieve the best possible alignment with Λ before calculating performance metrics.

#### 2.2.1 Scenario 1: Continuous Data

This scenario provides a baseline assessment against classical methods in a fully observed, continuous data setting.

Data are generated, as described in Section 2.2 with the observed data *Y* taken directly from the latent Gaussian vector, *Y* = *Z* and *α*_*d*_ = 0, *d* = 1, … *D*. We compare our method to principal component analysis (PCA) with Varimax rotation, Maximum Likelihood Factor Analysis (as implemented in the R package `factanal` (R Core Team, 2022)), and the sparse Bayesian factor analysis method of Ročková and George (2016). To ensure a fair comparison of loading matrix recovery, PCA and `factanal` are supplied with the true number of factors (*K* = 3). In contrast, our method and that of Ročková and George (2016) are tasked with inferring *K* directly from the data.

#### 2.2.2 Scenario 2: Binary Data

This scenario is designed to validate the model’s performance on fully observed binary data against appropriate competitors. We generate data for *N* = 200 individuals, as described in Section 2.2, where the observed binary data *Y*_*nd*_ ∈ {0, 1} is generated by setting 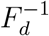 to a step function such that *p*(*Y*_*nd*_ = 1) = 0.25 +(*d* − 1)*/*22. Our approach is benchmarked against Nonnegative Matrix Factorisation (NMF), which encourages a sparse representation of the loading matrix as detailed in Kim and Park (2008), and the sparse Bayesian MIRT method of Li et al. (2023). As in the previous scenario, NMF is provided with the true *K* = 3, whereas our method and that of Li et al. (2023) infer the dimensionality from the data.

#### 2.2.3 Scenario 3: Mixed Type and Missing Data

This scenario showcases the model’s full capabilities in a realistic and challenging environment, characterised by mixed data types and structured missingness, for which the framework is developed. To construct a realistic setting, we base this scenario on a large clinical trial dataset with mixed outcome types and non random missingness, but work entirely in a simulated data framework. The reference dataset contains *N* = 8,023 individuals and eight variables of varying types (continuous, count, ordinal and binary). In the reference data, one variable exhibits approximately 20% missingness arising from study design. Further details on this reference dataset are given in Section 2.3.

Our simulation proceeds as follows. First, using the reference dataset, we fit a logistic regression model to estimate, for each individual, the probability that one of the variables is missing conditional on the remaining variables. We then generate complete datasets with mixed data types from the factor model with *D* = 8 observed variables, drawing latent Gaussian data and transforming each variable using the empirical cumulative distribution function *F*_*d*_ of the corresponding variable in the reference dataset. This preserves the marginal distributions and data types while keeping the dependence structure under our control. Finally, for each simulated dataset, we introduce missingness into the chosen variable according to the fitted logistic model, producing a structured missingness pattern that mimics the real data.

We evaluate our factor analysis performance under three conditions: full data, where the model is fitted to the complete generated data as an idealised benchmark; missing data, where the model is fitted to the incomplete datasets and must handle missingness directly; and complete cases, where all individuals with any missing observation are discarded prior to fitting. The true loading matrix is fixed and follows the sparse structure shown in fig. 2, with *D* = 8 observed variables.

**Figure 2:**
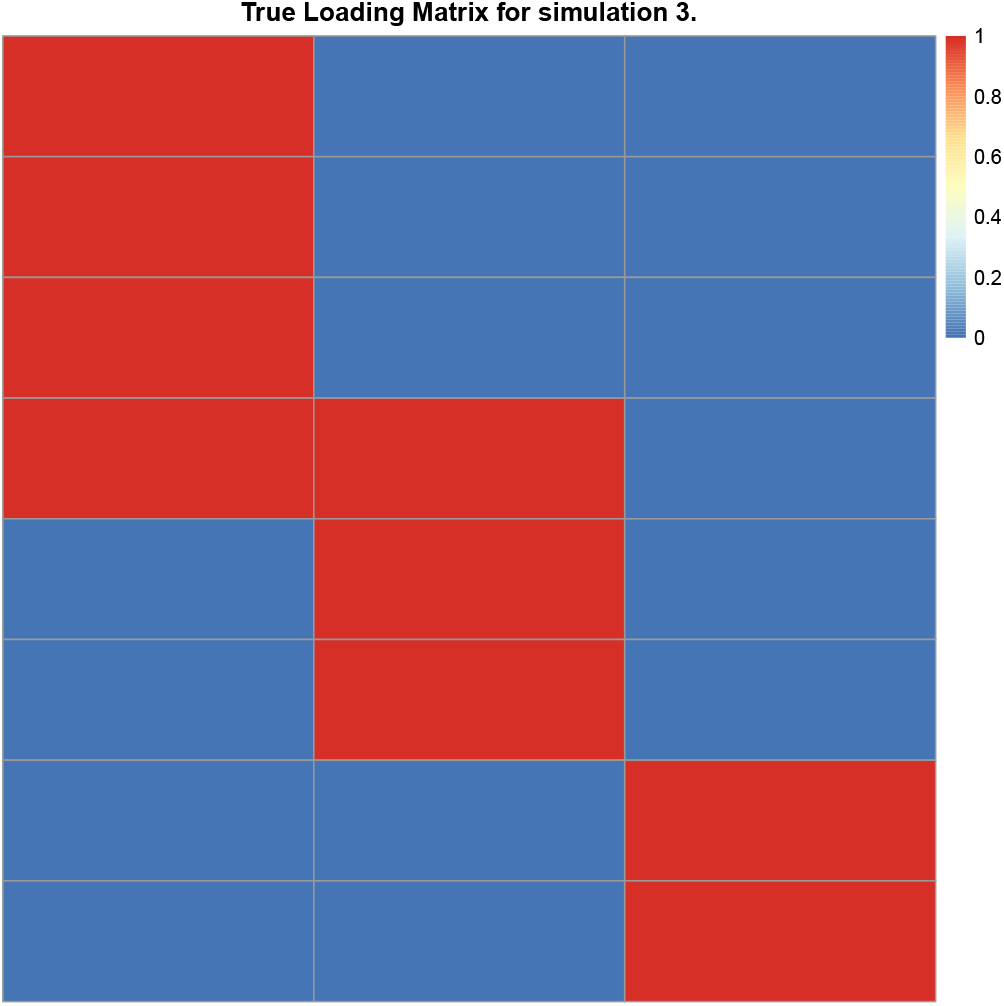
True 8 × K loading matrix used in simulation scenario 3.

### 2.3 Real Data

Multiple sclerosis (MS) is a debilitating disorder of the central nervous system, affecting approximately 2.9 million individuals worldwide (one in 3,000 people) (MS International Federation, 2023; Walton et al., 2020). The disease involves inflammation, neurodegeneration and destruction of axons, ultimately leading to severe physical and cognitive disability. Thousands of people living with MS have participated in global clinical trials or observational studies and provided clinical, biological and imaging data where multivariate modelling can provide better understanding of the disease which is essential for targeted drug development and appropriate patient treatment. The largest and most comprehensive of these clinical trial datasets is the Novartis-Oxford Multiple Sclerosis (NO.MS) dataset (Dahlke et al., 2021; Mallon et al., 2021), with up to 15 years of follow-up. To model such complex data, scalable methods are needed to reduce dimensionality in datasets with mixed modalities and to deal with structured missingness in a principled way. We evaluate our proposed model on this dataset to address two problems: first, identifying latent dimensions among MS clinical and radiological variables to characterise the disease to showcase our proposed method’s ability to handle different data modalities and structured missingness; and second, performing dimensionality reduction on structural MRIs to extract features for downstream analysis beyond traditional whole brain summary statistics.

The NO.MS data were collected across clinical trials conducted in accordance with the International Conference on Harmonization guidelines for Good Clinical Practice and the principles of the Declaration of Helsinki. All trial protocols were approved by an institutional review board or ethics committee and all patients or their legal representatives gave written informed consent before any trial-related procedures were performed. Data were de-identified following a risk-based approach as reported by Mallon et al. (2021).

#### 2.3.1 Multiple Sclerosis Dimensions

MS was originally classified according to consensus definitions based on an underlying survey (Lublin et al., 1996), and minimally revised to incorporate imaging features as disease modifiers (Lublin et al., 2014). The initial disease course descriptors were a pragmatic response to limited biological understanding and data availability, serving to provide a common clinical language to support trial design. Today, there is a significant need for data-driven, biologically informed classifications (Kuhlmann et al., 2023).

A key step towards a more data-driven characterisation of MS involves identifying latent dimensions from both clinical and radiological features, projecting them into a low dimensional continuous space. Ganjgahi et al. (2025) adopted this approach using eight commonly measured clinical trial variables. However, this approach had limitations: the method did not account for discrete data types, and was unable to accommodate missing data, which precluded the inclusion of variables not measured across all trials.

Here, we extend this line of work by analysing *N* = 8,023 individuals from NO.MS using nine baseline clinical and radiological variables comprising ordinal Expanded Disability Status Scale (EDSS), continuous assessments of physical function (timed 25-foot walk, 9-hole peg test), continuous cognitive assessments (symbol digit modalities test; SDMT), count-valued cognitive tests (paced auditory serial addition test; PASAT), Magnetic Resonance Imaging (MRI) derived measures (gadolinium-enhancing lesion counts, T2 lesion volume, normalised brain volume), and binary clinical relapse indicators. Crucially, the dataset exhibits substantial structured missingness arising from trial design: the constituent trials used different cognitive assessments, resulting in the PASAT being missing for 1,711 (21%) participants and the SDMT for 4,105 (51%) participants. This missingness arises directly from trial design rather than random omission. Unlike Ganjgahi et al. (2025), our framework handles mixed data types and structured missingness directly, which allows us to include both the PASAT and the SDMT despite these not being measured in all trials.

For identifying latent dimensions of MS, we analyse the nine variables mentioned above, using DPE with a cooling schedule *v*_0_ ∈ {1, 0.8, 0.4, 0.2, 0.1, 0.05} and both rotation and scale parameter expansions.

Each DPE stage runs for 101 EM iterations with 100 samples per E-step. The model is fitted from 10 random initialisations, all of which recover a similar factor structure. We report results from a single run. The simulation scenario described in Section 2.2.3 was informed by these data, using all variables except the SDMT.

#### 2.3.2 Spatial Modes of Lesion Co-occurrence

The accumulated brain damage from past and ongoing neuroinflammation appears as hyperintense areas in T2-weighted MRI, known as T2 lesions. The distribution of T2 lesions in time and space is important for MS diagnosis, and the accumulation of such lesions in terms of count and volume is used for disease monitoring and prognostication. However, lesion count and volume are only two summary statistics extracted from brain images containing thousands of volume elements (voxels), and they lack spatial information about lesion shape, location and pattern of co-occurrence. Such summary statistics correlate only weakly with clinical outcomes (Barkhof, 2002). Directly modelling high dimensional, binary lesion maps can capture spatial variation (Altermatt et al., 2018), but voxelwise mass univariate approaches are simplistic, treating voxels independently and failing to model spatial correlations. Consequently, the predictive value of each voxel is weak, and fully Bayesian methods that could model spatial dependencies do not scale to data of this dimensionality.

Hence, there is a need to extract clinically meaningful summaries from high dimensional lesion maps that retain spatial information and the pattern of lesion co-occurrence. Such summaries can be used for downstream analysis, such as predicting disability worsening, and may provide prognostic value beyond simpler metrics that discard lesion location.

We apply our proposed framework to compress baseline binary lesion masks from *N* = 2,093 people living with relapsing-remitting MS (RRMS) from the FREEDOMS (NCT00289978) and FREEDOMS II (NCT00355134) trials in NO.MS. The lesion masks were obtained using an automated segmentation approach (Sun et al., 2025) and nonlinearly registered to MNI space. We compress these into a small number of interpretable spatial summaries. The columns of the estimated loading matrix identify spatial patterns of co-occurring lesions, which we term Modes of lesion Co-occurrence (MoCos), while the factor scores quantify each individual’s expression of each MoCo. Together, these scores provide a low dimensional representation of each individual’s lesion pattern, suitable for use as imaging-derived covariates in downstream analysis. The model is initialised with a maximum of *K* = 100 latent dimensions under the IBP stick-breaking truncation from a random initialisation and fitted using DPE (*v*_0_ ∈ {10^−1^, 10^−2^, 10^−3^}, *v*_1_ = 10) with the scale parameter expansion of Murray et al. (2013). The first DPE stage uses a convergence tolerance of 0.01 on the maximum absolute loading change, and for subsequent stages, parameter differences are monitored until they plateau. We assess clinical relevance by fitting Cox proportional hazards models for time to 3-month confirmed disability worsening (we refer the reader to Lublin et al., 2022, for details regarding this definition), including MoCo scores as covariates and controlling for age, sex, active treatment status, disease duration, number of relapses in the year prior to trial entry, EDSS, gadolinium-enhancing lesion count, and normalised brain volume.

## 3 Results

We evaluate the proposed framework through simulation, then apply it to the NO.MS dataset to address two questions central to MS research: First, can we identify clinically meaningful latent dimensions of disease status from incomplete data of mixed types? Second, can we compress high dimensional lesion maps into spatial summaries that provide prognostic value beyond traditional imaging metrics?

### 3.1 Simulations

Simulation scenario 1 assesses performance on continuous, fully observed data, comparing our method against PCA, `factanal` (R Core Team, 2022), and Ročková and George (2016).

The results are summarised in table 1. Our method correctly identifies the latent dimensionality in 75.8% of replications and recovers the true sparsity structure in 61.4%. The only other method capable of inferring both, Ročková and George (2016), achieves substantially lower rates of 13.3% and 12.7% respectively. In terms of loading matrix recovery, our method attains an MSE of 0.0023, lower than factanal (0.0054) and Ročková and George (2016) (0.023), and more than an order of magnitude lower than PCA (0.036).

**Table 1:** Performance comparison on continuous data. MIDFA is compared against standard factor analysis (`factanal`), PCA, and the sparse Bayesian method of Ročková and George (2016). ‘NA’ indicates metrics not applicable for methods that do not yield sparse loadings or infer the number of latent dimensions.

| Method | Struct. Rec. Rate | Correct $K$ Rate | $\Lambda$ MSE (SE) |
| --- | --- | --- | --- |
| MIDFA | 0.614 (0.0218) | 0.758 (0.0192) | 0.002262 (0.0003341) |
| factanal ($K$ known) | NA | NA | 0.005426 (0.0000658) |
| PCA ($K$ known) | NA | NA | 0.036045 (0.0003928) |
| Ročková and George (2016) | 0.127 (0.0149) | 0.133 (0.0152) | 0.022501 (0.0008230) |

The NA entries for PCA and `factanal` reflect that these methods do not produce sparse loading matrices and therefore structure recovery is not defined. Likewise, because both methods are given the true *K* as input, a “correct *K* rate” is not meaningful.

Simulation scenario 2 evaluates performance on binary data, comparing our method against NMF and the sparse MIRT model of Li et al. (2023).

The results are presented in table 2. Our method correctly identifies the latent dimensionality in 82.2% of replications and recovers the true sparsity structure in 65.8%. The only other method capable of inferring both, Li et al. (2023), achieves lower rates of 54.2% and 42.8% respectively, despite using twice as many DPE stages. In terms of loading matrix recovery, our method attains an MSE of 0.0047, compared to 0.0058 for Li et al. (2023) and 0.036 for NMF.

**Table 2:** Performance comparison on binary data. MIDFA is benchmarked against nonnegative matrix factorisation and the sparse MIRT model of Li et al. (2023). ‘NA’ indicates metrics not applicable for methods that do not yield sparse loadings or infer the number of latent dimensions.

| Method | Struct. Rec. Rate | Correct $K$ Rate | $\Lambda$ MSE (SE) |
| --- | --- | --- | --- |
| MIDFA | 0.658 (0.0212) | 0.822 (0.0171) | 0.004658 (0.0006003) |
| NMF ($K$ known) | NA | NA | 0.035505 (0.0001073) |
| Li et al. (2023) | 0.428 (0.0221) | 0.542 (0.0223) | 0.005791 (0.0003085) |

As in scenario 1, the NA entries for NMF appear because it does not produce sparse loading matrices and is supplied with the true *K*.

Simulation scenario 3 evaluates performance under mixed data types with structured missingness, the setting most relevant to real clinical studies. The marginal distributions and missingness patterns were derived from the NO.MS data to mimic realistic conditions. We compare three conditions: full data (benchmark), structured missingness (one covariate missing for a subset of individuals), and complete case analysis.

The results are summarised in table 3. Under structured missingness, our method correctly identifies the latent dimensionality in 99.2% of replications and recovers the true sparsity structure in 72.8%. These rates are within two standard errors of the full data benchmark (99.4% and 71.8% respectively) and exceed complete case analysis (98.0% and 67.0%), which discards individuals with partially observed data, losing information that remains recoverable under our model. In terms of loading matrix recovery, the MSE under structured missingness is 0.0047, compared to 0.0049 for the full data benchmark and 0.0076 for complete case analysis.

**Table 3:** Performance comparison under mixed data and structured missingness. Results are shown for the full data benchmark, the setting with simulated structured missingness, and a complete case analysis in which individuals with any missing value are removed.

| Analysis Condition | Struct. Rec. Rate | Correct $K$ Rate | $\Lambda$ MSE (SE) |
| --- | --- | --- | --- |
| Full Data (benchmark) | 0.718 (0.0201) | 0.994 (0.0035) | 0.004860 (0.0006802) |
| Structured Missingness | 0.728 (0.0199) | 0.992 (0.0040) | 0.004682 (0.0006551) |
| Complete Case Analysis | 0.670 (0.0210) | 0.980 (0.0063) | 0.007629 (0.0008501) |

### 3.2 Latent Dimensions to Characterise Multiple Sclerosis

We first applied the proposed model to NO.MS to determine whether a small number of interpretable latent dimensions could summarise the key clinical and radiological features of MS. The model inferred the presence of *K* = 5 latent dimensions. The corresponding sparse loading matrix is shown in table 4, where blank entries indicate loadings that are more likely to belong to the spike component and are therefore treated as zero. The resulting sparsity facilitates a clear interpretation of each latent dimension.

**Table 4:** Estimated loading matrix, Λ, for latent dimensions of MS. Blank entries indicate loadings assigned to the spike component.

|  | Physical disability | Cognitive function | Asymptomatic activity | Relapse | Brain damage |
| --- | --- | --- | --- | --- | --- |
| EDSS | 1.43 |  |  |  |  |
| Timed 25-Foot Walk | 1.24 |  |  |  |  |
| 9-Hole Peg Test | 0.90 |  |  |  |  |
| Paced Auditory Serial Addition Test |  | 1.02 |  |  |  |
| Symbol Digit Modalities Test | −0.71 | 1.13 |  |  |  |
| T2 lesion volume |  |  | 1.12 |  | −0.86 |
| Normalised brain volume |  |  |  |  | 0.95 |
| Number of Gd-enhancing T1 lesions |  |  | 1.28 |  |  |
| Relapse status |  |  |  | 1.06 |  |

The first latent dimension, physical disability, is characterised by strong positive loadings on EDSS, the Timed 25-Foot Walk and the 9-Hole Peg Test, all objective assessments of physical function. The second dimension reflects cognitive function, driven by large loadings from the SDMT and the PASAT. The SDMT additionally loads negatively on the physical disability dimension. The third dimension corresponds to asymptomatic activity, driven by the number of gadolinium-enhancing T1 lesions and T2 lesion volume, consistent with asymptomatic radiological disease activity. The fourth dimension is associated solely with relapse status and therefore reflects whether a patient is experiencing acute new or worsening neurological symptoms. The fifth dimension corresponds to brain damage, combining normalised brain volume and T2 lesion volume, and captures overall radiological disease burden.

Four of these dimensions (physical disability, brain damage, relapse and asymptomatic activity) closely align with those reported in the recent reclassification analysis of Ganjgahi et al. (2025). Notably, our estimated loadings are about two to three times larger in magnitude than those of Ganjgahi et al. (2025). This difference is likely due to two aspects of our method: the principled handling of missing data, which allows the full sample to be retained rather than restricting analysis to complete cases; and the Gaussian copula formulation, which respects the mixed data types of the observed variables. Neither of these features is present in the approach of Ganjgahi et al. (2025). In addition, our model uncovers a distinct cognitive function dimension, separating cognitive function from brain damage, which loaded together as the same dimension in Ganjgahi et al. (2025). This dimension becomes identifiable because the model explicitly accounts for the structured, trial level missingness in cognitive assessments, which is substantial in the NO.MS dataset. By modelling this missingness mechanism directly rather than discarding incomplete observations, the method is able to recover a clinically coherent latent dimension that would otherwise be obscured. These results demonstrate that the framework successfully addresses the first problem, recovering interpretable latent dimensions of MS from incomplete data of mixed types.

### 3.3 Spatial Modes of Lesion Co-occurrence

We next assess whether the framework can compress high dimensional lesion data into clinically meaningful summaries. Applied to binary lesion masks from *N* = 2,093 individuals in NO.MS, the model compresses 51,595 voxels into *K* = 100 Modes of lesion Co-occurrence (MoCos). The spike and slab prior induces sparsity in the loading matrix, resulting in spatially localised MoCos that capture distinct patterns of lesion co-occurrence. The inferred MoCos are spatially localised despite no spatial prior, indicating that the model identifies common lesion distributions purely from the data structure (see fig. 3). This contrasts with principal component analysis, which yields dense, non localised components that are difficult to interpret (fig. 4).

**Figure 3:**
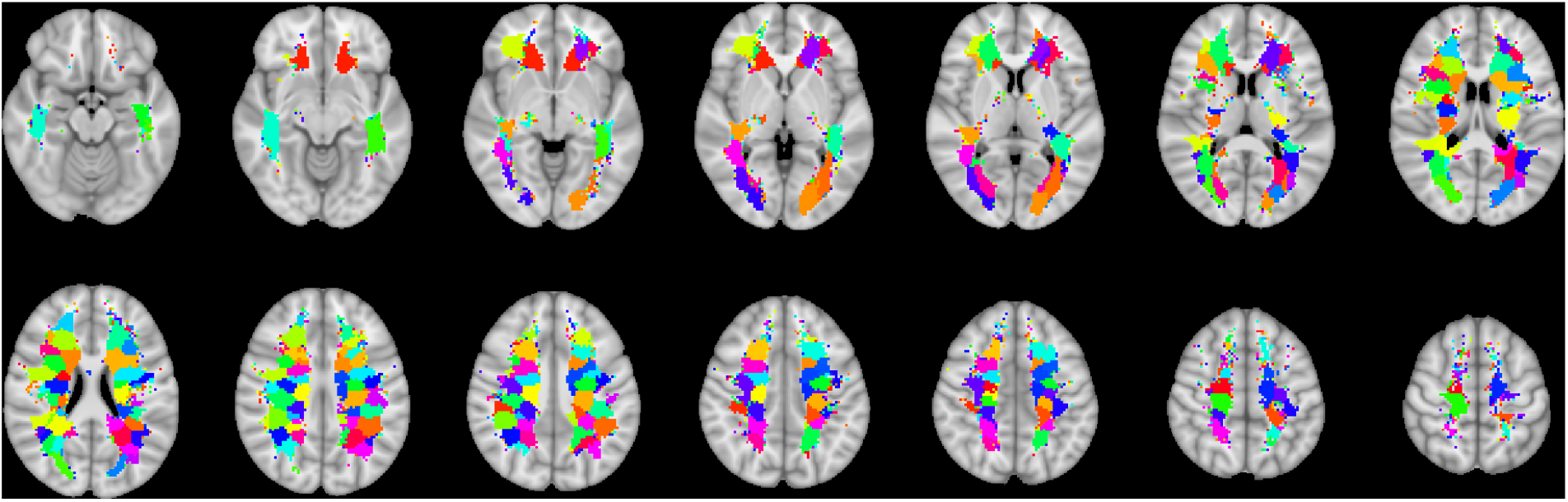
Spatial maps (loading matrix Λ) corresponding to the 100 MoCos discovered by the model. Each MoCo reflects a distinct spatial mode of lesion co-occurrence.

**Figure 4:**
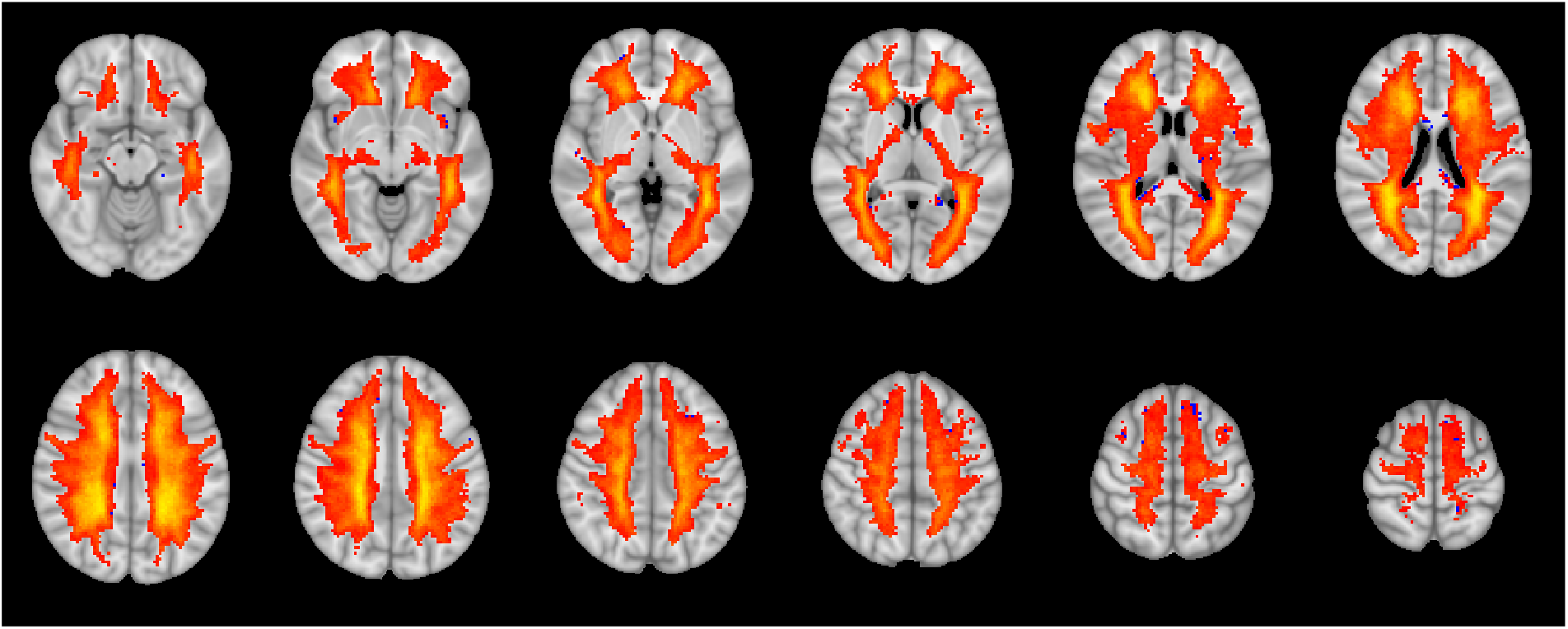
Spatial map of the first principal component. PCA produces dense, non localised components, illustrating the limitations of traditional linear methods for lesion data.

To assess prognostic value, we include the MoCo scores as covariates in a Cox model for time to confirmed disability worsening, adjusting for baseline covariates. Two MoCos show significant associations: MoCo 48 (*p* = 0.0058) and MoCo 76 (*p* = 0.0083). These findings show that specific MoCos are predictive of 3-month confirmed disability worsening (fig. 5). By contrast, total T2 lesion volume is not predictive in a Cox model with the same baseline covariates (*p* = 0.5554), underscoring the limitations of traditional global summary metrics.

**Figure 5:**
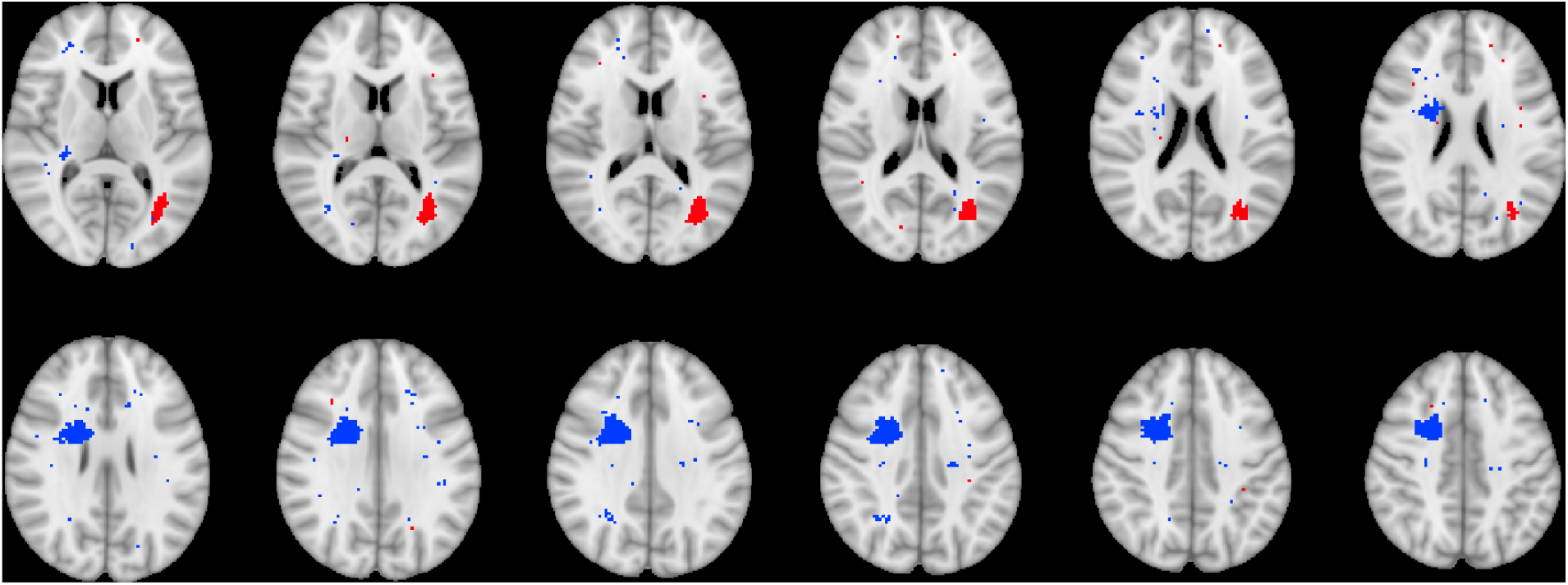
Spatial maps for the two MoCos whose scores are significantly associated with confirmed disability worsening in a Cox model: MoCo 48 (blue) and MoCo 76 (red).

Taken together, these findings show that the proposed model uncovers sparse MoCos that are predictive of 3-month confirmed disability worsening.

## 4 Discussion

In this paper, we introduced a scalable Bayesian factor analysis framework designed to address the challenges presented by modern, large scale biomedical datasets. These datasets involve high dimensionality, mixed data types and substantial structured missingness. Existing methods typically address some of these difficulties in isolation, but few provide a unified approach capable of handling all of them simultaneously. Our framework combines these capabilities within a single model, and, when latent factors correspond to meaningful biological or clinical patterns, the sparse structure encouraged by the prior can make these patterns easier to interpret. At the same time, the framework remains applicable to more modestly sized datasets, uncovering coherent, clinically meaningful latent dimensions from the mixed data type NO.MS data. This combination of flexibility and scalability makes the framework well suited to a broad range of biomedical applications.

Methodologically, the framework combines several components. It (i) maps marginal distributions to a latent Gaussian scale via a semiparametric copula, (ii) fits a sparse latent factor model using a continuous spike and slab prior, (iii) infers the number of latent dimensions nonparametrically using an Indian Buffet Process prior, and (iv) fits the model using a scalable, EM like scheme that naturally accommodates structured missingness. This modular construction allows the dependence structure, sparsity pattern and latent dimensionality to be learned jointly from the data while keeping inference computationally tractable.

Our simulation studies further demonstrate the strengths of the proposed framework across a range of data types and missingness patterns. In continuous data, the model achieves the highest structure recovery rate and lowest loading matrix error among all methods considered. Similar performance is seen in binary data. The most challenging setting, involving mixed data types and structured missingness, highlights an important practical advantage of the approach. In this scenario, the marginal distributions and variable types were matched to those observed in the NO.MS data, and the missingness pattern was generated to approximate the structured, trial level mechanism present in the real data, where specific variables are systematically unobserved for subsets of individuals due to study design. When fitted directly to these incomplete datasets, the model maintains strong performance comparable to the full data benchmark, and exceeds complete case analysis. Taken together, these results demonstrate that the proposed framework is robust to realistic missingness patterns and retains high accuracy in settings that mirror the complexities of large scale biomedical studies.

The application to the NO.MS dataset highlights the practical utility of the framework. For identifying latent dimensions of MS from incomplete data of mixed types, the model recovers five dimensions that provide a compact summary of disease status. Four of these dimensions, reflecting physical disability, brain damage, relapse and asymptomatic activity, align closely with those reported in recent reclassification work (Ganjgahi et al., 2025), while an additional cognitive function dimension emerges because the model explicitly accounts for the trial level missingness in the cognitive assessments. This illustrates the value of modelling structured missingness directly, allowing the framework to recover latent structure that would otherwise be obscured when incomplete variables are discarded.

For compressing high dimensional lesion maps into sparse spatial summaries, the framework reduces 51,595 voxels to 100 MoCos. Two MoCos are predictive of 3-month confirmed disability worsening after adjustment for standard clinical and imaging covariates. These findings indicate that the MoCos capture spatial lesion patterns that are clinically meaningful and prognostically informative, offering a substantially richer summary of lesion distribution than conventional global imaging metrics such as total T2 lesion volume, which was not significantly associated with 3-month confirmed disability worsening.

Despite its strong performance, our framework has several limitations. The dependence structure is based on a Gaussian copula, which assumes latent linear relationships and may not capture more complex, non linear dependencies. The use of the extended rank likelihood, while computationally convenient and robust, is an approximation to the full likelihood and may result in some loss of information. For fully continuous data, as noted by Murray et al. (2013), one can avoid this approximation by transforming the observed variables to a latent Gaussian scale using the empirical CDFs and then treating these transformed variables as fixed. Finally, while the EM algorithm provides a scalable solution for finding a posterior mode, it can still be computationally intensive for extremely large datasets and, like all EM based methods, it is guaranteed only to converge to a local optimum.

Several avenues for future research are apparent. The modularity of the framework allows for the substitution of more flexible copula structures and alternative priors. For instance, incorporating spatial priors, such as those proposed by Menacher et al. (2024), could further enhance the interpretability of the neuroimaging factors by explicitly encoding the known spatial structure of the brain. While EM provides a scalable solution for MAP estimation, developing a fully Bayesian MCMC based inference scheme, perhaps building on recent advances in high dimensional MCMC, could provide richer posterior uncertainty quantification for smaller scale problems. Finally, the successful application of this framework to the NO.MS dataset suggests its potential utility for a wide range of other large scale biomedical studies facing similar analytical challenges.

## Data and Code Availability

The real data used in this study are from the Novartis-Oxford Multiple Sclerosis (NO.MS) dataset, which is not publicly available. The reader is able to request the raw data (anonymised) and related documents by connecting to CSDR (https://www.clinicalstudydatarequest.com) and signing a data-sharing agreement with Novartis, with requests reviewed and approved by an independent review panel of CSDR. The code used in this study is available at https://github.com/george-hutchings/factor-analysis-paper-code.

## Author Contributions

GH: Conceptualization, Formal analysis, Investigation, Methodology, Software, Validation, Visualization, Writing - original draft, Writing - review & editing. PS: Software, Writing - review & editing. CD: Writing - review & editing. LG: Resources, Writing - review & editing. EF: Resources, Writing - review & editing. TEN: Project administration, Supervision, Writing - review & editing. CH: Project administration, Supervision. DAH: Resources, Writing - review & editing. HG: Conceptualization, Data curation, Methodology, Project administration, Supervision, Writing - review & editing.

## Supplementary Material

### EM Algorithm Derivations

The Expectation-Maximisation (EM) algorithm is an iterative method for finding maximum a posteriori (MAP) estimates of parameters when the model involves latent variables or missing data (Dempster et al., 1977). Let *Y* denote the observed data, *Ƶ* = {*Z, η, w*} the collection of all latent variables, and Θ = {Λ, *α, θ*} the parameters. For cases where the E-step is intractable, this procedure can be adapted (Ročková & George, 2016), as below:

1. **E-Step (Sampling):** Given the current parameter estimates Θ^(*t*)^, draw *M* samples of the latent variables from their conditional posterior distribution: 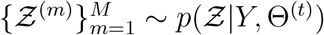.
2. **E-Step (Expectation):** Approximate the expected complete-data log-posterior using the Monte Carlo samples:

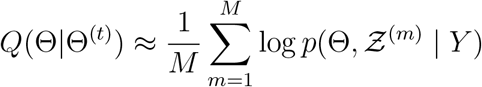
3. **M-Step (Maximisation):** Find the parameter estimates that maximise this function:

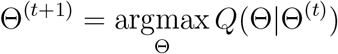

These steps are repeated until convergence. Below we derive the conditional posteriors for the sampling step and the update equations for the maximisation step.

#### E-Step

The E-step requires calculating expectations with respect to *p*(*Z, η, w* | *Y*, Θ^(*t*)^).

Notice that the right hand side separates into a product of two factors: one involving only (*Z, η*) and the other involving only *w*,

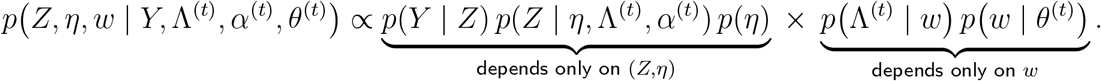

Hence the conditional posterior factorises, and the *Q*-function separates (up to additive constants). In particular, since the second factor depends only on *w* given Θ^(*t*)^ and not on *Y*, the expectation with respect to *w* is conditioned on Θ^(*t*)^ alone:

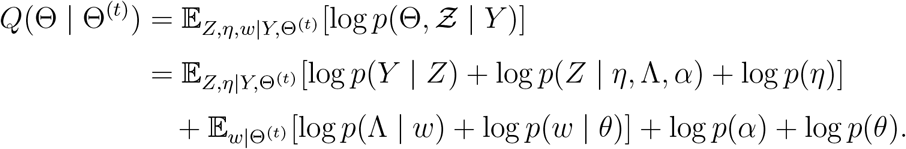

Due to this separation we can split the E step into two stages, calculating expectations with respect to *η, Z* and calculating expectations with respect to *w*.

#### Expectations with respect to *η* **and** *Z*

We calculate expectations with respect to *η* and *Z* through approximation with samples, hence we wish to sample *Z, η*.

Given all other parameters, the full conditional for *Z* factorises across features *d* = 1, …, *D*. Let *Z*_•*d*_ = (*Z*_1*d*_, …, *Z*_*Nd*_)^⊤^, and *Y*_•*d*_ = (*Y*_1*d*_, …, *Y*_*Nd*_)^⊤^, the full conditional is:

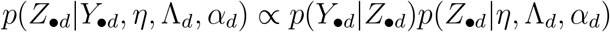

The term *p*(*Y*_•*d*_|*Z*_•*d*_) is an indicator function, 𝕀(*Z*_•*d*_ ∈ 𝒟(*Y*_•*d*_)), that enforces the rank constraints, and *p*(*Z*_•*d*_|*Y*_•*d*_, *η*, Λ_*d*_, *α*_*d*_) = 𝒩 (*Z*_•*d*_|*η*Λ_*d*_ + *α*_*d*_1_*N*_, *I*_*N*_). This results in a truncated multivariate normal posterior. We can sample each element *Z*_*nd*_ univariately:

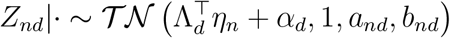

where the truncation bounds [*a*_*nd*_, *b*_*nd*_] are determined as described in the main text, following eq. (15). For missing *Y*_*nd*_, the distribution is a non truncated Normal.

For the factor scores, *p*(*η*|Λ, *α, Y*) factorises across individuals *n* = 1, …, *N*. For each individual’s factor score *η*_*n*_:

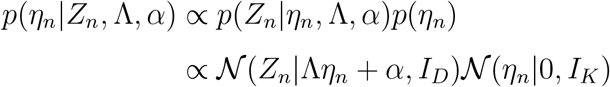

The log of this is quadratic in *η*_*n*_, implying a Gaussian distribution 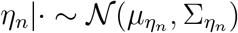, where completing the square yields:

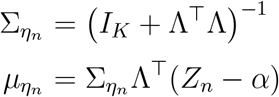

#### Expectation with respect to *w*

The posterior for the binary indicators *w*_*dk*_ factorises for each loading. The conditional probability for *w*_*dk*_ being in the slab (*w*_*dk*_ = 1) is:

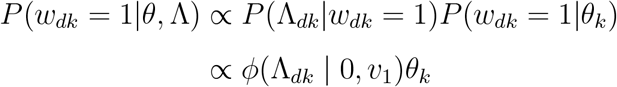

Normalising gives the required expectation:

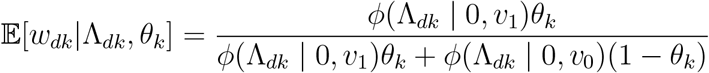

where *ϕ*(· | 0, *v*) denotes the univariate Gaussian density with mean zero and variance *v*.

#### M-Step

In the M-step, we maximise the *Q*-function. Throughout this section, 𝔼[·] denotes expectation under the E-step posterior *p*(*Ƶ* | *Y*, Θ^(*t*)^) unless otherwise indicated. Given that these expectations can be evaluated as described in the E-step, the *Q*-function can be broken down as follows (up to additive constants):

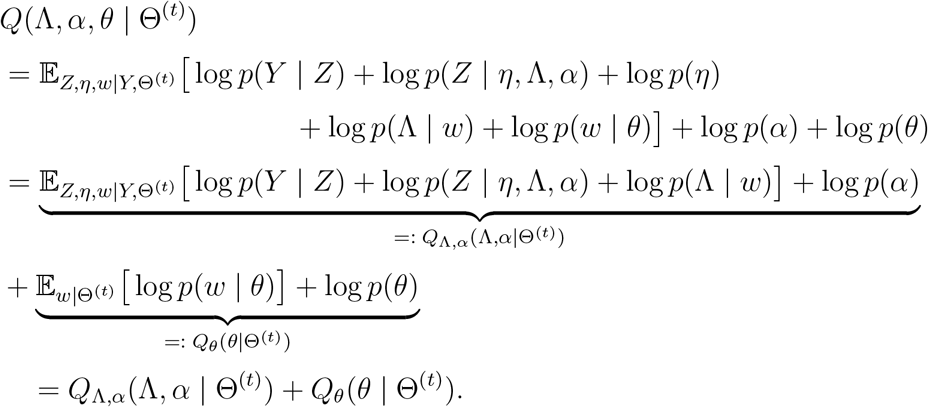

These derivations demonstrate that the *Q*-function separates into distinct components, allowing the parameter updates to be carried out independently: (Λ, *α*) are updated by maximising *Q*_Λ,*α*_(Λ, *α* | Θ^(*t*)^), and *θ* is updated by maximising *Q*_*θ*_(*θ* | Θ^(*t*)^).

#### Update for Loadings Λ

Restrict attention to the terms of the *Q*-function that depend on Λ_*d*_. Dropping constants that do not involve Λ_*d*_ we have

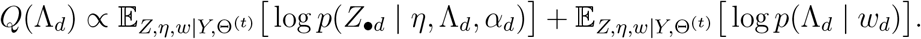

Substituting the Gaussian forms and expanding yields a quadratic form in Λ_*d*_:

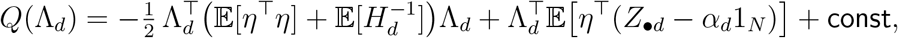

where *H*_*d*_ = diag{(1 − *w*_*dk*_)*v*_0_ + *w*_*dk*_*v*_1_} and the expectations are taken under *p*(*Z, η, w* | *Y*, Θ^(*t*)^).

Completing the square shows that exp(*Q*(Λ_*d*_)) is proportional to a multivariate Gaussian density with precision

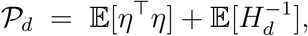

and mean

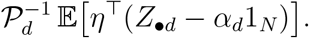

Hence the M-step maximiser of *Q* with respect to Λ_*d*_ is

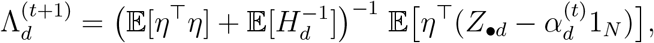

Note that (*H*_*d*_)_*kk*_ = (1 − *w*_*dk*_)*v*_0_ + *w*_*dk*_*v*_1_, so 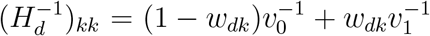. Taking expectations with respect to the E-step posterior of *w*_*dk*_ yields

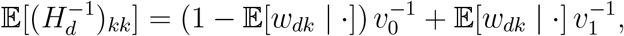

and hence 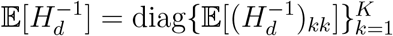.

#### Update for intercepts *α*

The intercepts *α*_*d*_ decouple across features. Restricting the *Q*-function to terms that depend on *α*_*d*_ and dropping constants gives

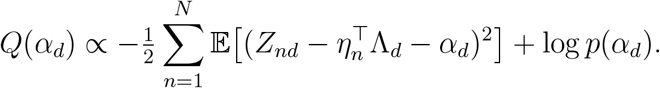

For a Gaussian prior *α*_*d*_ ~ 𝒩 (0, 1), completing the square and differentiating yields the closed form update:

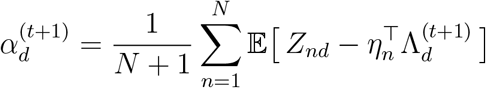

#### Monte Carlo approximation of the updates

The updates for Λ_*d*_ and *α*_*d*_ derived above involve expectations of functions of (*Z, η*) under the E-step posterior, which are not available in closed form. We approximate them by averaging over the *M* Gibbs samples 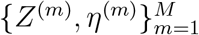 drawn in the E-step:

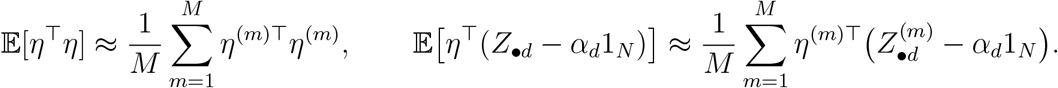

The expectation 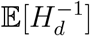 depends only on *w*, whose posterior inclusion probabilities 𝔼[*w*_*dk*_] are available in closed form, so no sampling approximation is needed for this term.

Substituting these approximations into the exact updates yields the computable forms:

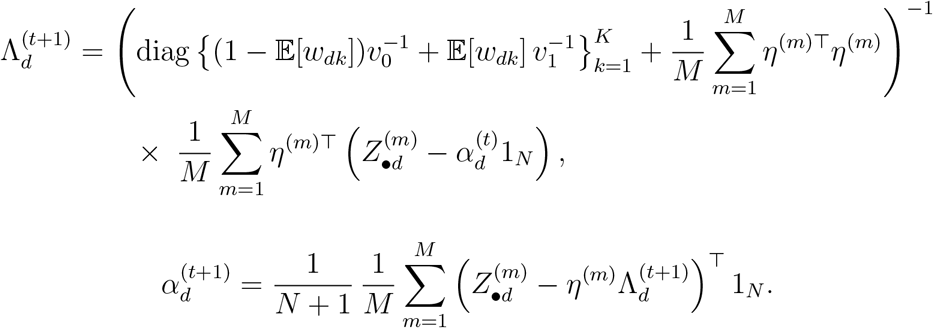

#### Update for IBP parameters *θ*

The objective function for *θ* is derived from the priors on *w* and the stick breaking construction for *u*_*k*_:

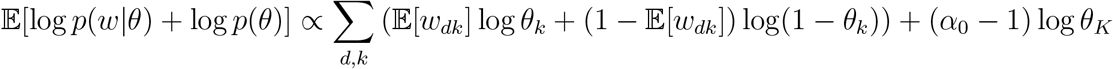

This function is maximised with respect to *θ* subject to the ordering constraint 1 *> θ*_1_ > · · · *> θ*_*K*_ > 0 using a numerical optimisation routine.

## References

Albert, J. H., & Chib, S. (1993). Bayesian analysis of binary and polychotomous response data. Journal of the American statistical Association, 88 (422), 669–679.

Altermatt, A., Gaetano, L., Magon, S., Häring, D. A., Tomic, D., Wuerfel, J., Radue, E.-W., Kappos, L., & Sprenger, T. (2018). Clinical Correlations of Brain Lesion Location in Multiple Sclerosis: Voxel-Based Analysis of a Large Clinical Trial Dataset. Brain Topography, 31 (5), 886–894. 10.1007/s10548-018-0652-9

Barkhof, F. (2002). The clinico-radiological paradox in multiple sclerosis revisited. Current opinion in neurology, 15 (3), 239–45.

Casey, B. J., Cannonier, T., Conley, M. I., Cohen, A. O., Barch, D. M., Heitzeg, M. M., Soules, M. E., Teslovich, T., Dellarco, D. V., Garavan, H., et al. (2018). The adolescent brain cognitive development (abcd) study: Imaging acquisition across 21 sites. Developmental cognitive neuroscience, 32, 43–54.

Comon, P. (1994). Independent component analysis, a new concept? Signal processing, 36 (3), 287–314.

D. Hoff P. (2007). Extending the rank likelihood for semiparametric copula estimation.

Dahlke, F., Arnold, D. L., Aarden, P., Ganjgahi, H., Häring, D. A., Čuklina, J., Nichols, T. E., Gardiner, S., Bermel, R., & Wiendl, H. (2021). Characterisation of ms phenotypes across the age span using a novel data set integrating 34 clinical trials (no. ms cohort): Age is a key contributor to presentation. Multiple Sclerosis Journal, 27 (13), 2062–2076.

Dempster, A. P., Laird, N. M., & Rubin, D. B. (1977). Maximum likelihood from incomplete data via the em algorithm. Journal of the royal statistical society: series B (methodological), 39 (1), 1–22.

Feldman, J., & Kowal, D. R. (2022). Bayesian data synthesis and the utility-risk trade-off for mixed epidemiological data. The Annals of Applied Statistics, 16 (4), 2577–2602.

Ganjgahi, H., Häring, D. A., Aarden, P., Graham, G., Sun, Y., Gardiner, S., Su, W., Berge, C., Bischof, A., Fisher, E., et al. (2025). Ai-driven reclassification of multiple sclerosis progression. Nature Medicine, 1–11.

George, E. I., & McCulloch, R. E. (1993). Variable selection via gibbs sampling. Journal of the American Statistical Association, 88 (423), 881–889.

Ghahramani, Z., & Griffiths, T. (2005). Infinite latent feature models and the indian buffet process. Advances in neural information processing systems, 18.

Hastie, T., Tibshirani, R., Friedman, J., et al. (2009). The elements of statistical learning.

Kaiser, H. F. (1958). The varimax criterion for analytic rotation in factor analysis. Psychometrika, 23 (3), 187–200.

Kim, H., & Park, H. (2008). Nonnegative matrix factorization based on alternating nonnegativity constrained least squares and active set method. SIAM journal on matrix analysis and applications, 30 (2), 713–730.

Kuhlmann, T., Moccia, M., Coetzee, T., Cohen, J. A., Correale, J., Graves, J., Marrie, R. A., Montalban, X., Yong, V. W., Thompson, A. J., et al. (2023). Multiple sclerosis progression: Time for a new mechanism-driven framework. The Lancet Neurology, 22 (1), 78–88.

Le Roux, B., & Rouanet, H. (2010). Multiple correspondence analysis (Vol. 163). Sage.

Li, J., Gibbons, R., & Rockova, V. (2023). Sparse bayesian multidimensional item response theory. arXiv preprint arXiv:2310.17820.

Lublin, F. D., Häring, D. A., Ganjgahi, H., Ocampo, A., Hatami, F., Čuklina, J., Aarden, P., Dahlke, F., Arnold, D. L., Wiendl, H., et al. (2022). How patients with multiple sclerosis acquire disability. Brain, 145 (9), 3147–3161.

Lublin, F. D., Reingold, S. C., Cohen, J. A., Cutter, G. R., Sørensen, P. S., Thompson, A. J., Wolinsky, J. S., Balcer, L. J., Banwell, B., Barkhof, F., et al. (2014). Defining the clinical course of multiple sclerosis: The 2013 revisions. Neurology, 83 (3), 278–286.

Lublin, F. D., Reingold, S. C., & on Clinical Trials of New Agents in Multiple Sclerosis*, N. M. S. S. (A. C. (1996). Defining the clinical course of multiple sclerosis: Results of an international survey. Neurology, 46 (4), 907–911.

Mallon, A.-M., Häring, D. A., Dahlke, F., Aarden, P., Afyouni, S., Delbarre, D., El Emam, K., Ganjgahi, H., Gardiner, S., Kwok, C. H., et al. (2021). Advancing data science in drug development through an innovative computational framework for data sharing and statistical analysis. BMC medical research methodology, 21 (1), 250.

Menacher, A., Nichols, T. E., Holmes, C., & Ganjgahi, H. (2024). Bayesian lesion estimation with a structured spike-and-slab prior. Journal of the American Statistical Association, 119 (545), 66–80.

Mitchell, T. J., & Beauchamp, J. J. (1988). Bayesian variable selection in linear regression. Journal of the american statistical association, 83 (404), 1023–1032.

MS International Federation. (2023). Atlas of MS [Data updated 2023. Accessed: 2025-12-07].

Murray, J. S., Dunson, D. B., Carin, L., & Lucas, J. E. (2013). Bayesian gaussian copula factor models for mixed data. Journal of the American Statistical Association, 108 (502), 656–665.

Pearson, K. (1901). Liii. on lines and planes of closest fit to systems of points in space. The London, Edinburgh, and Dublin Philosophical Magazine and Journal of Science, 2 (11), 559–572.

Quinn, K. M. (2004). Bayesian factor analysis for mixed ordinal and continuous responses. Political Analysis, 12 (4), 338–353.

R Core Team. (2022). R: A language and environment for statistical computing. R Foundation for Statistical Computing. Vienna, Austria. https://www.R-project.org/

Ročková, V., & George, E. I. (2016). Fast bayesian factor analysis via automatic rotations to sparsity. Journal of the American Statistical Association, 111 (516), 1608–1622.

Rubin, D. B. (1987). Multiple imputation for nonresponse in surveys. Wiley.

Sklar, M. (1959). Fonctions de repartition an dimensions et leurs marges. Publ. inst. statist. univ. Paris, 8, 229–231.

Spearman, C. (1961). “general intelligence” objectively determined and measured.

Sterne, J. A., White, I. R., Carlin, J. B., Spratt, M., Royston, P., Kenward, M. G., Wood, A. M., & Carpenter, J. R. (2009). Multiple imputation for missing data in epidemiological and clinical research: Potential and pitfalls. Bmj, 338.

Sudlow, C., Gallacher, J., Allen, N., Beral, V., Burton, P., Danesh, J., Downey, P., Elliott, P., Green, J., Landray, M., et al. (2015). Uk biobank: An open access resource for identifying the causes of a wide range of complex diseases of middle and old age. PLoS medicine, 12 (3), e1001779.

Sun, Y., Gaetano, L., Fisher, E., Haring, D., Aarden, P., Hallan-Nickell, S., Pehrs-Jurvillier, A.-C., Bendfeldt, K., Brugger, C., Bermel, R., et al. (2025). An automated deep learning segmentation tool for accurate and generalizable ms lesion quantification. MULTIPLE SCLEROSIS JOURNAL, 31 (3), 261–262.

Teh, Y. W., Grür, D., & Ghahramani, Z. (2007). Stick-breaking construction for the indian buffet process. Artificial intelligence and statistics, 556–563.

Van Essen, D. C., Ugurbil, K., Auerbach, E., Barch, D., Behrens, T. E., Bucholz, R., Chang, A., Chen, L., Corbetta, M., Curtiss, S. W., et al. (2012). The human connectome project: A data acquisition perspective. Neuroimage, 62 (4), 2222–2231.

Walton, C., King, R., Rechtman, L., Kaye, W., Leray, E., Marrie, R. A., Robertson, N., La Rocca, N., Uitdehaag, B., van Der Mei, I., et al. (2020). Rising prevalence of multiple sclerosis worldwide: Insights from the atlas of ms. Multiple Sclerosis Journal, 26 (14), 1816–1821.

